# Cancer Cell Line Encyclopedia Data Suggest that Ligands for ERBB Family Receptors May Drive *BRAF*-WT Melanomas

**DOI:** 10.64898/2026.03.16.712185

**Authors:** Ella Wilson, Andrew Conway, David J. Riese

## Abstract

Cutaneous skin melanomas with wild-type BRAF alleles (“*BRAF*-WT melanomas”) remain relatively difficult to treat, even though they typically possess driver mutations in a *RAS* gene or *NF1*. For example, these tumors respond relatively poorly to combinations of MEK and BRAF inhibitors, and their response to ICIs is muted compared to the response of *BRAF-*mutant melanomas. *ERBB2* and *ERBB4*, which encode receptor tyrosine kinase genes, are necessary and sufficient for the proliferation of multiple *BRAF*-WT melanoma cell lines. Consequently, we have postulated that ERBB4-ERBB2 heterodimerization drives *BRAF-*WT melanomas. This mechanism is consistent with the observation that elevated *ERBB4* transcription or *ERBB4* mutations are found in a significant fraction of *BRAF*-WT melanoma tumor samples. Moreover, a subset of *ERBB4* mutations found in *BRAF*-WT melanoma samples increases proliferation in a B*RAF*-WT melanoma cell line. Because the elevated *ERBB4* transcription observed in *BRAF-* WT melanomas is typically insufficient to cause ligand-independent ERBB4 signaling, we have postulated that ligands for ERBB family receptors drive the elevated ERBB4-ERBB2 heterodimerization responsible for the proliferation of BRAF-WT melanoma cell lines. We have explored this hypothesis by analyzing data found in the Broad Institute’s Cancer Cell Line Encyclopedia. These data suggest that some EGF family hormones are required for the proliferation of *BRAF*-WT melanoma cell lines. Likewise, the G_α11_/G_αq_ pathway, which can stimulate cleavage and maturation of EGF family hormones, is also required for the proliferation of *BRAF*-WT melanoma cell lines. Thus, these data suggest additional therapeutic targets in *BRAF*-WT melanomas. Moreover, because many uveal (ocular) melanomas possess elevated G_α11_/G_αq_ signaling, these data suggest that ligand stimulation of ERBB receptor signaling may contribute to uveal melanomagenesis or progression.

## II. Introduction

There have been recent significant improvements in the treatment of cutaneous skin melanomas (“melanomas”) that have gain-of-function *BRAF* mutations. Many of these tumors respond to the combination of MEK and BRAF inhibitors. Many of these tumors also respond to immune checkpoint inhibitors (ICIs) [1]. Melanomas with wild-type BRAF alleles (“*BRAF*-WT melanomas”) remain relatively difficult to treat, even though they typically possess driver mutations in a *RAS* gene or *NF1*. For example, these tumors respond relatively poorly to combinations of MEK and BRAF inhibitors, and their response to ICIs is muted compared to the response of *BRAF-*mutant melanomas [1, 2].

Thus, there is substantial interest in identifying actionable targets for therapeutic intervention in *BRAF*-WT melanomas. Our group has investigated ERBB4, a receptor tyrosine kinase closely related to the Epidermal Growth Factor Receptor (EGFR/ERBB2), ERBB2/HER2, and ERBB3/HER4. Detectable *ERBB4* transcription is observed in approximately 10% of *BRAF*-WT melanoma samples [3], suggesting that elevated ERBB4 signaling drives *BRAF*-WT melanomas. Similarly, approximately 40 different candidate gain-of-function *ERBB4* mutants are found in *BRAF*-WT melanoma samples [4]. The population of *BRAF*-WT melanoma samples that exhibit “elevated” *ERBB4* transcription does not substantially overlap with the population in which *ERBB4* is mutated [4], suggesting that *ERBB4* transcription and *ERBB4* mutation are independent drivers of *BRAF*-WT melanomas. Ectopic overexpression of wild-type *ERBB4* stimulates the proliferation of four *BRAF*-WT melanoma cell lines [3] (IPC-298 [5], MEL-JUSO [6], MeWo [7], and SK-MEL-2 [8]). Similarly, expression of a dominant-negative *ERBB4* mutant inhibits the proliferation of these cell lines [3].

Some *ERBB4* mutants identified in *BRAF*-WT melanoma samples exhibit a gain-of-function phenotype in MEL-JUSO cells. Surprisingly, the synthetic *ERBB4* Q646C mutant, which encodes a constitutively homodimerized and active ERBB4 protein, inhibits the proliferation of MEL-JUSO cells [4]. This apparent conundrum is largely resolved by the observation that *ERBB2* is both sufficient and necessary for the proliferation of IPC-298, MEL-JUSO, and MeWo cells [9]; in these cell lines, ERBB4-ERBB2 heterodimers (rather than ERBB4-ERBB4 homodimers) apparently drive proliferation.

In *BRAF*-WT melanoma samples that exhibit “elevated” *ERBB4* transcription, the absolute level of *ERBB4* transcription does not appear to be sufficient to enable ligand-independent ERBB signaling [3], which appears to require approximately 300 thousand to 1 million receptors per cell [10]. Therefore, we have postulated that the elevated ERBB4 signaling in *BRAF*-WT melanoma cells is largely ligand-dependent. We note that ERBB receptor heterodimerization could enable ERBB4, ERBB3, or EGFR ligands to stimulate signaling by ERBB4-ERBB2 heterodimers [11].

To begin exploring this hypothesis, we have examined the Broad Institute’s Cancer Cell Line Encyclopedia (CCLE) [12] for evidence of ERBB ligand activity in four *ERBB4*-dependent, *BRAF*-WT melanoma cell lines (IPC-298, MEL-JUSO, MeWo, and SK-MEL-2). These data suggest multiple potential mechanisms for elevated ERBB ligand activity in these cells.

## III. Results and Discussion

### A. Mutation, Gene Transcription, RNAi, and CRISPR Data

We downloaded DepMap Public 25Q3 data from the Broad Institute’s CCLE on December 11, 2025. These data included mutation calls, gene transcription, effects of RNA interference (RNAi) on cell proliferation, and effects of CRISPR gene knockout on cell proliferation. The transcription, RNAi, and CRISPR data are all reported as log2 values. We have highlighted RNAi and CRISPR data that are -0.2 or less or 0.2 or greater. These highlighted variations represent a change in proliferation (relative to controls) of approximately 15%.

### B. Cell Lines and Genes Studied in These Analyses

We have studied two male (MeWo and SK-MEL-2 and two female (IPC-298 and MEL-JUSO) *ERBB4*-dependent, *BRAF*-WT melanoma cell lines. We used the A-375, G-361, SK-MEL-1, and SK-MEL-28 BRAF-mutant melanoma cell lines as one set of controls. We used the MM28, MP41, and MP46 uveal (ocular) melanoma cell lines as another set of controls. The spreadsheet of these cell lines and the other cell lines found in the Broad Institute’s CCLE is provided (**Supplemental File 1**).

We studied the expression and activity of several genes. They include genes encoding components of the RAS signaling pathway (*HRAS, KRAS, NRAS, NF1*), genes encoding members of the Epidermal Growth Factor (EGF) family (*AREG, BTC, EGF, EPGN, EREG, HBEGF, NRG1, NRG2, NRG3, NRG4, TGFA*), genes encoding components of the PI3 kinase pathway (*PIK3CA, PTEN*), and genes encoding components of the G_α11_/Gαq pathway (*GNAQ, GNA11, PLCB4*). A spreadsheet of these genes is provided (**Supplemental File 2**)

### C. Genes Encoding Components of the RAS Signaling Pathway Are Required

Most *BRAF*-WT melanomas harbor a gain-of-function mutation in a *RAS* allele or a loss-of-function mutation in *NF1* alleles, leading to elevated RAS signaling [13]. We and others have reported that the IPC-298, MEL-JUSO, MeWo, and SK-MEL-2 *BRAF*-WT melanoma cell lines also possess these genetic changes. The Broad Institute’s CCLE data are consistent with these previous reports and indicate that RAS signaling is required for the proliferation of the IPC-298, MEL-JUSO, and SK-MEL-2 cell lines. Because CRISPR data are not available for the MeWo cell line, it is unclear whether RAS Signaling is required for its proliferation (**Table 1**).

**Table 1.**

| Cell Line | Gene Name | Mutations | Transcription | RNAi Effect | CRISPR Dependency | CRISPR Effect |
| --- | --- | --- | --- | --- | --- | --- |
| IPC298<br>ACH-000915 | HRAS | NR | 5.021 | -0.111 | 0.076 | -0.136 |
|  | KRAS | NR | 4.052 | -0.096 | 0.037 | -0.063 |
|  | NRAS | Q61L | 6.764 | -1.955 | 1.000 | -1.877 |
|  | NF1 | NR | 3.965 | -0.060 | 0.005 | 0.118 |
| MELJUSO<br>ACH-000881 | HRAS | G13D | 6.495 | -0.693 | 0.996 | -1.351 |
|  | KRAS | NR | 3.746 | -0.245 | 0.820 | -0.672 |
|  | NRAS | Q61L | 5.213 | -0.485 | 1.000 | -1.931 |
|  | NF1 | L1779P | 3.727 | -0.109 | 0.014 | 0.006 |
| MEWO<br>ACH-000987 | HRAS | NR | 5.985 | -0.058 | ND | ND |
|  | KRAS | NR | 5.805 | -0.114 | ND | ND |
|  | NRAS | NR | 5.407 | -0.106 | ND | ND |
|  | NF1 | Q1336Ter | 2.937 | -0.114 | ND | ND |
| SKMEL2<br>ACH-001190 | HRAS | NR | 4.690 | 0.034 | 0.070 | -0.140 |
|  | KRAS | NR | 5.284 | 0.086 | 0.059 | -0.119 |
|  | NRAS | Q61L | 7.121 | -1.419 | 1.000 | -2.907 |
|  | NF1 | NR | 4.734 | 0.021 | 0.085 | -0.167 |

NR: No value reported ND: No data

### D. Genes Encoding Components of the PI3K Signaling Pathway Are Required

Approximately 25% of *BRAF*-WT melanoma samples possess elevated *PIK3CA* transcription, a gain-of-function *PIK3CA* mutation, reduced *PTEN* transcription, or a loss-of-function *PTEN* mutation [4]. These data suggest that elevated PI3 kinase (PI3K) signaling drives the proliferation of *BRAF*-WT melanoma cell lines. However, these potential drivers of increased PI3K signaling are inversely correlated with candidate *ERBB4* driver mutations in *BRAF*-WT melanomas, suggesting that elevated signaling by ERBB4-ERBB2 heterodimers in *BRAF*-WT melanomas stimulates PI3K signaling independently of these potential causes.

None of the four *BRAF*-WT melanoma cell lines possesses a gain-of-function *PIK3CA* mutation or a loss-of-function *PTEN* mutation. However, CRISPR data indicate that the *PIK3CA* gene is required for the proliferation of the IPC-298, MEL-JUSO, and SK-MEL-2 cell lines. CRISPR data also indicate that *PTEN* gene disruption stimulates the proliferation of these cell lines (**Table 2**). These data indicate that PI3K signaling is both necessary and sufficient for the proliferation of the IPC-298, MEL-JUSO, and SK-MEL-2 cell lines.

**Table 2.**

| Cell Line | Gene Name | Mutations | Transcription | RNAi Effect | CRISPR Dependency | CRISPR Effect |
| --- | --- | --- | --- | --- | --- | --- |
| IPC298<br>ACH-000915 | PIK3CA | NR | 3.412 | -0.173 | 0.246 | -0.282 |
|  | PTEN | NR | 4.791 | 0.026 | 0.000 | 1.052 |
| MELJUSO<br>ACH-000881 | PIK3CA | NR | 3.008 | -0.121 | 0.253 | -0.318 |
|  | PTEN | NR | 5.178 | 0.021 | 0.000 | 0.270 |
| MEWO<br>ACH-000987 | PIK3CA | NR | 3.181 | 0.088 | ND | ND |
|  | PTEN | NR | 4.632 | -0.061 | ND | ND |
| SKMEL2<br>ACH-001190 | PIK3CA | NR | 4.327 | -0.027 | 0.369 | -0.411 |
|  | PTEN | NR | 4.813 | -0.079 | 0.000 | 0.594 |

NR: No value reported ND: No data

Unfortunately, CRISPR data were not available for the MeWo cell line.

### E. Genes that Encode Some Ligands for ERBB Receptors May Be Required

Many of the genes that encode EGF family hormones exhibit at least modest transcription in the BRAF-WT melanoma cell lines. Exceptions include the betacellulin (BTC) gene in the IPC-298 and SK-MEL-2 cell lines; the epigen (*EPG*) gene in the IPC-298, MEL-JUSO, and MeWo cell lines; the epiregulin (*EREG*) gene in the IPC-298, MEL-JUSO, MeWo, and SK-MEL-2 cell lines; and the neuregulin-1 (NRG1) gene in the IPC-298 cell line (**Table 3**). The *NRG1* gene is mutated (G119E) in the MeWo cell line, whereas the TGFA gene is mutated (S26R) in the SK-MEL-2 cell line (**Table 3**). These expression and mutation data suggest that the activity of EGF family hormones may drive the proliferation of *BRAF*-WT melanomas.

**Table 3.**
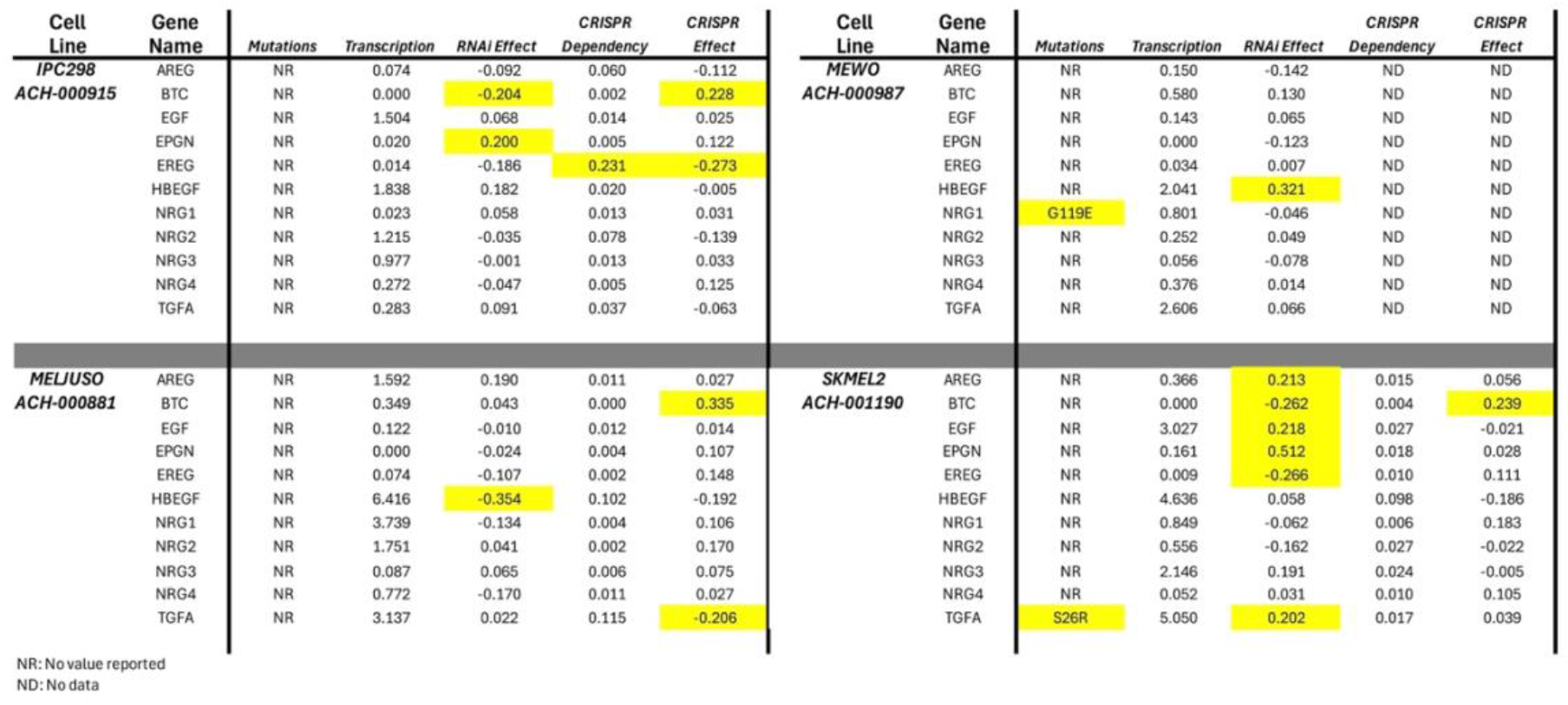

RNAi targeting the *BTC* gene reduces IPC-298 proliferation by a modest amount, as does RNAi targeting *HBEGF* in MEL-JUSO cells (**Table 3**). Using CRISPR to disrupt the *EREG* gene causes a modest decrease in IPC-298 proliferation, as does CRISPR against the Transforming Growth Factor alpha (*TGFA*) gene in MEL-JUSO cells (**Table 3**). The modest effects of these RNAi and CRISPR approaches and the failure of other RNAi and CRISPR approaches suggest that multiple EGF family hormones may be responsible for driving the proliferation of each *BRAF*-WT melanoma cell line. Thus, identifying mechanisms by which multiple EGF family hormones can be simultaneously regulated may lead to effective strategies for blocking ligand-induced signaling by ERBB4-ERBB2 heterodimers.

### E. The G_α11_/G_αq_ Signaling Pathway May Regulate the Activity of EGF Family Hormones

EGF family hormones are expressed as transmembrane precursor proteins. Cleavage of these precursors releases the mature, soluble form of the protein, which can bind EGFR, ERBB3, or ERBB4. Members of the matrix metalloproteinase (MMP) or ADAM (a disintegrin and metalloproteinase) are responsible for this cleavage, and the activity of these proteases can be regulated by G-protein coupled receptors (GPCRs). For example, GPCRs coupled to G_α11_ or G_αq_ can stimulate the activity of the SRC tyrosine kinase via activation of phospholipase C beta 4 (PLCβ4). SRC stimulation of MMP or ADAM activity results in cleavage of the transmembrane precursor form of an EGF family hormone, releasing the mature hormone into the extracellular milieu. This enables hormone binding to ERBB receptors and prototypical ligand-induced ERBB receptor signaling [14-30].

Mutations in the *PLCB4* gene are found in the MEL-JUSO, MeWo, and SK-MEL-2 cell lines, while a mutation in the *GNA11* gene is found in the IPC-298 cell line (**Table 4**). CRISPR disruption of the *GNAQ* gene inhibits the proliferation of the IPC-298, MEL-JUSO, and SK-MEL-2 cell lines, whereas RNAi against the *PLCB4* gene inhibits the proliferation of the IPC-298 cell line (Table 4). These data suggest that the G_α11_/G_αq_ signaling pathway is required for the proliferation of *BRAF-*WT melanoma cell lines, perhaps by stimulating the cleavage and activity of EGF family hormones.

**Table 4.**
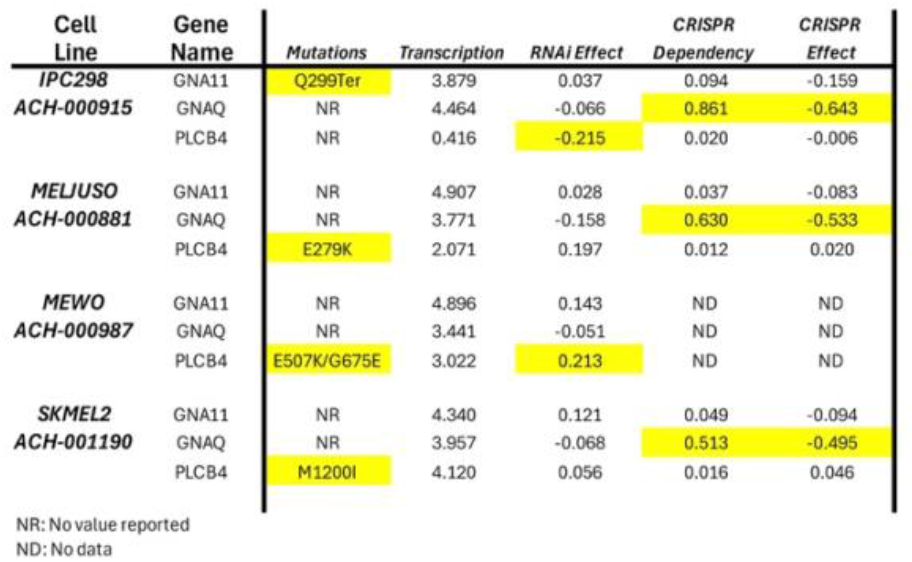

Uveal (ocular) melanoma (UM) is a tumor arising from melanocytes in the uveal tissues (iris, ciliary body, and choroid). The most common genetic alterations in UM involve mutations in the *GNAQ* or *GNA11* genes [31-35], resulting in constitutive canonical GPCR/G_αq_/G_α11_ signaling through loss of G_αq_ or G_α11_ GTPase activity [35]. Uveal melanomas have a very high (∼50%) incidence of metastasis and mortality, even in patients with no evidence of residual disease following removal of the affected eye [33-42]. Thus, there is considerable interest in targeted approaches to treating uveal melanoma micrometastases and preventing their expansion. Our data suggest that EGF family hormones or ERBB receptors may be suitable targets for such studies.

## V. Supplemental Information

All supplemental files have been compressed into a ZIP archive, named <SupplementalFiles.zip>.

Supplemental File 1: <RelevantCellLines.xlsx>

Spreadsheet of the cell lines studied in this manuscript, as well as the other cell lines found in the Broad Institute’s CCLE.

Supplemental File 2: <GenesOfInterest1.xlsx>

Spreadsheet of the genes studied in this manuscript.

Supplemental File 3: <AggregatedTables.xlsx>

Spreadsheet of the data reported in this manuscript.

## Notes

### Competing Interest Statement

The authors have declared no competing interest.

### Funding

- National Institutes of Health: R15CA280767-01A1

